# Antimetastatic dsRNA mimics identified by live imaging of pathogenic neolymphangiogenesis

**DOI:** 10.1101/2019.12.26.887943

**Authors:** David Olmeda, Daniela Cerezo-Wallis, Tonantzin G. Calvo, Direna Alonso, Estela Cañón, Nuria Ibarz, Javier Muñoz, Sagrario Ortega, María S. Soengas

**Affiliations:** Melanoma Laboratory, Molecular Oncology Programme, Spanish National Cancer Research Centre (CNIO), Madrid, Spain; Proteomics Unit, Biotechnology Programme, Spanish National Cancer Research Centre (CNIO), Madrid, Spain; Transgenic Mice Unit, Biotechnology Programme (CNIO)

**Keywords:** GEMM-melanoma models, Non-invasive tumor imaging, Neo-lymphangiogenesis, Tumor relapse, MIDKINE, pre-metastatic niche, dsRNA-nanoplexes

## Abstract

The crosstalk between cancer cells and the lymphatic vasculature has long been proposed to define competency for metastasis. Nevertheless, the discovery of selective blockers of lymphovascular niches has been compromised by the paucity of experimental systems for whole-body analyses of tumor progression. Here we present immunocompetent and immunodeficient mouse models for live imaging of melanoma-induced neolymphangiogenesis (driven by Vegfr3) as a cost-effective platform for drug screening *in vivo*. Spatio-temporal analyses in autochthonous melanomas and patient-derived xenografts identified double stranded RNA mimics (dsRNA nanoplexes) as potent repressors of lymphangiogenesis and metastasis. Mechanistically, dsRNA nanoplexes were found to suppress lymphangiogenic drivers in both tumor cells and their associated lymphatic vasculature (via MIDKINE and Vegfr3, respectively). This dual inhibitory action, driven by type I interferon, was not shared by FDA-approved antimelanoma treatments or by lymphangiogenic blockers in clinical testing. These results underscore the power of Vegfr3-lymphoreporters for pharmacological testing in otherwise aggressive cancers.

**RELEVANCE:** Although tumor-induced lymphangiogenesis has long been associated with metastasis, selective targeting of this process has been compromised by the paucity of experimental platforms for whole-body imaging of tumor progression and drug response. Here we present animal models engineered for spatio-temporal analyses of neolymphangiogenesis in clinically relevant autochthonous melanomas and patient-derived xenografts, and identify a unique action of double stranded-RNA nanoplexes as potent repressors of lymphatic dissemination and metastatic relapse.

## INTRODUCTION

Clinical intervention in the cancer field has been revolutionized by the identification of (epi)genetic alterations in tumor cells as the basis for rational drug design^1,2^. Prime example of this success is malignant melanoma, where BRAF mutations have led to the generation of inhibitors with unprecedented efficacy^3,4^. Unfortunately, responses to BRAF blockers and other genetically-targeted agents are characteristically transient due to a plethora of mechanisms of resistance^5–9^. Immune checkpoint blockers (e.g. anti-PD1, -PDL1 or -CTLA4) are providing unprecedented response rates, particularly in combination with targeted therapies^10–14^. Nevertheless, toxicities may be limiting^10,11,15^, and median progression-free survival remains below 3 years^11,16^. The field is therefore, actively seeking for alternative and more effective treatments. Rather unexplored from a therapeutic perspective (in melanoma and other tumor types) is the lymphatic system. The rationale for this approach stems from the observation that expansion of the tumoral lymphatic vasculature (neo-lymphangiogenesis) is one of the earliest events in the dissemination of a variety of aggressive neoplasms^17–19^. Antagonists of the interaction of lymphangiogenic ligands (i.e. VEGFC/D) with their receptors (VEGFR family members) are under evaluation in multiple diseases^17,20,21^. However, whether sustained therapeutic responses can be achieved with these agents has yet to be demonstrated^19^.

The development of anti-lymphangiogenic compounds has been compromised by the lack of suitable models for live imaging of distant metastatic niches^22^. Among lymphatic biomarkers, Vegfr3 represents a conceptual platform for drug screening as this gene is selectively enriched under pathological situations (e.g. cancer), while being highly downregulated in normal adult endothelial cells^23,24^. We have reported melanoma “Met-Alert lymphoreporter” mice to monitor the endogenous transcription of *Vegfr3*^*25*^. These animals take advantage of a targeted *knock in* strategy to express a EGFP-Luciferase cassette coupled to the expression of *Vegfr3*^*26*^. The Vegfr3^Luc^ models have revealed long range-acting mechanisms of neolymphangiogenesis and melanoma metastasis driven by the heparin binding factor Midkine (MDK)^25^. Here we evaluate the potential of the *Vegfr3*^*Luc*^-lymphoreporters for *in vivo* discovery of anti-cancer agents. To this end, *Vegfr3*^*Luc*^ immunodeficient mice were tested for whole-body imaging of tumors induced by human melanoma cells and patient-derived xenografts. In parallel, immunocompetent *Vegfr3*^*Luc*^-mice were exploited for the assessment of autochthonous melanocytic lesions driven by oncogenic *Braf* and *Pten* loss. These models were analyzed in two scenarios that recapitulate main clinical needs: (i) established melanomas, and (ii) relapsed tumors after surgical excision of primary lesions. This strategy identified a sustained therapeutic action of dsRNA-based nanoparticles in both these settings. While this study focused on melanoma, our results support the *Vegfr3*^*Luc*^ mice as a versatile strategy for pharmacological analyses of aggressive cancers where the lymphatic system plays a key role in metastasis.

## RESULTS

### Acute blockade of melanoma-induced linfovascular niches by dsRNA mimics

Given the impact of neo-lymphangiogenesis on the conditioning and colonization of distant metastatic niches in melanoma^25^, we hypothesized that “lymphoreporter” mice could serve as tractable platform for preclinical studies of anticancer agents. To this end, we used genetically engineered mice models (GEMM) where tumor-associated lymphangiogenesis is driven by autochthonous melanomas induced by conditional activation of oncogenic *Braf*^V600E^ in the context of *Pten* loss (Fig 1a). This was achieved by crosses of immunocompetent *Vegfr3*^*Luc*^ with the melanocyte-specific *Tyr:CreERT2;Braf*^*V600E*^;*Pten*^*flox/flox*^ strain^27^ (Supplementary Fig. S1a; see Supplementary Materials for additional detail). *Vegfr3*^*Luc*^ mice were also crossed into nude (*nu/nu*) mice to generate suitable hosts for whole-body imaging of tumors generated by human melanoma cell lines (Fig. 1b), or human patient-derived xenografts (PDX, Fig. 1c). As described before^25^, the intrinsically low luciferase emission in the *Vegfr3*^*Luc*^ mice is greatly induced as melanomas progress to metastasis (see examples for basal vs tumor-induced bioluminiscence in Fig. 1a, left panels). Animals were randomized for treatment when Vegfr3-driven luciferase was detected in a systemic manner, namely at proximal and distant sites (inguinal, axillary and brachial lymph nodes), as well as visceral sites (see Figs. 1a-c). The BRAF inhibitor vemurafenib was used as representative example of FDA-approved melanoma treatments actively used in the clinic^28^. As shown in Fig. 1a, *Vegfr3*^*Luc*^;*Tyr:CreERT2;Braf*^*V600E*^;*Pten*^*flox/flox*^ mice were indeed responsive to this compound (50 mg/kg, 7 doses per week x 3 weeks), with the expected reduction in tumor growth and consequently, in the emission of Vegfr3-luciferase (see also Fig. S1b for macroscopic analyses and for histological assessment of tyrosinase-related protein 2 (Trp2) and phospho-ERK as readouts for melanocytic cells and MAPK/ERK inhibition, respectively). However, the therapeutic effects of vemurafenib were short-lived, as melanomas ultimately regained growth (Fig. 1d). Therefore, we screened for more potent blockers of neolymphangiogenesis and metastatic dissemination. Dacarbazine/DTIC was selected as a negative control in this analysis, as an example of an alkylating agent used for decades for the treatment of metastatic melanomas (most recently for NRAS-mutated cases) but now nearly discontinued for its minimal efficacy^29^. DITC (80 mg/kg, 5 doses/week, 2 weeks) had no significant effect on luciferase emission at primary tumors or distant sites (see results for NRAS-melanomas in Figs. S1c-d). This maintained Vegfr3-luc signal was consistent with a minimal impact of DTIC on melanoma growth (Fig. 1e).

**Figure 1.**
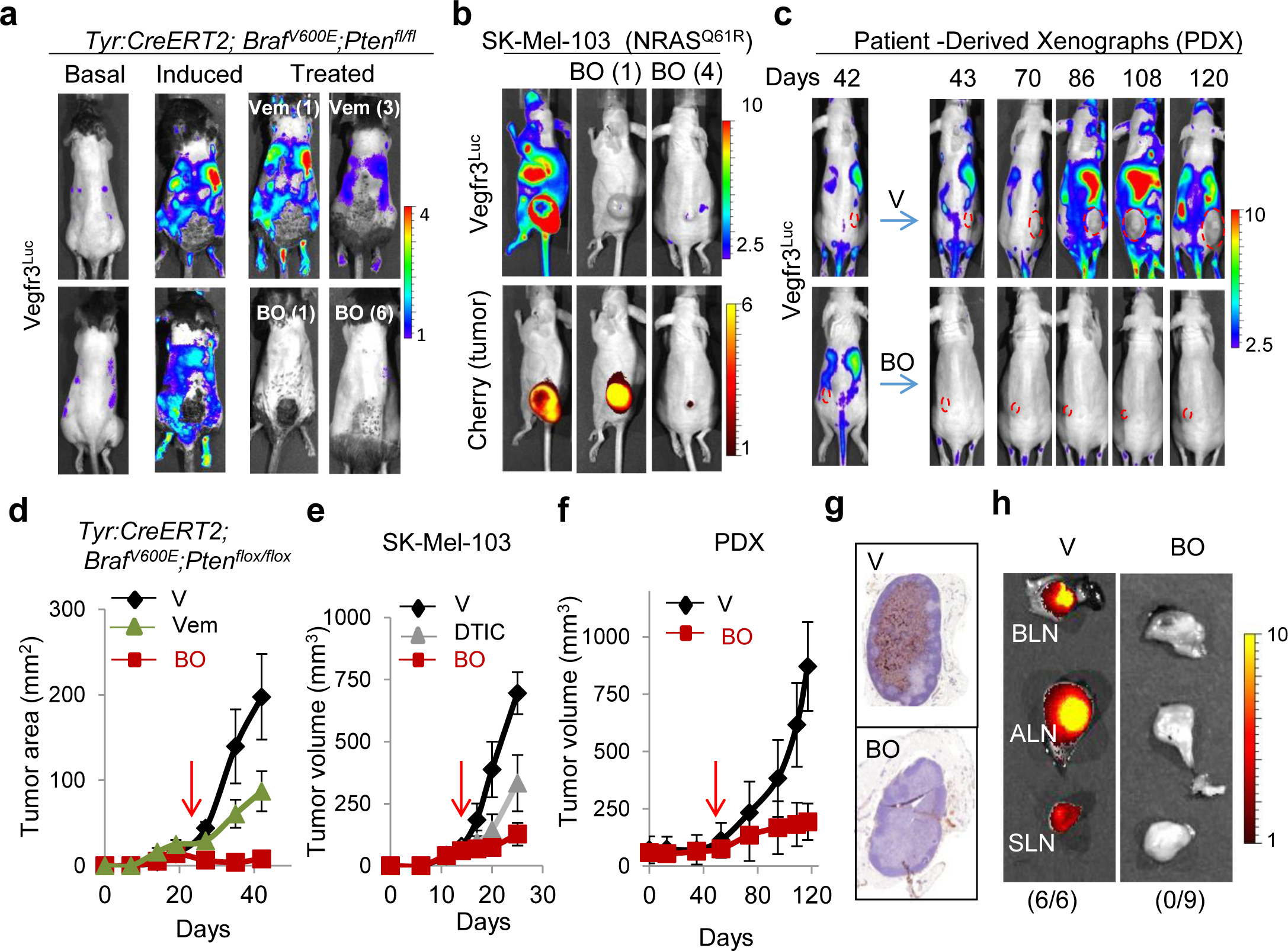
Identification of anti-lymphangiogenic compounds in *Vegfr3*^*Luc*^ GEMM mice. **(a)** Luciferase-based imaging of drug response in *Vegfr3*^*Luc*^;*Tyr:CreERT2*;*Braf*^*V600E*^;*Pten*^*flox/flox*^; mice. Panels labeled as “basal” and “induced” correspond to bioluminescence of animals prior and 5 weeks after administration of tamoxifen (4H-Tamoxifen 5 mM, topical administration, 3 consecutive days) for the induction of melanomas. Animals were then treated with one or six doses of BO-110 (BO, 0.8 mg/kg, twice per week, three weeks) or with one or 21 doses of Vemurafenib (50 mg/Kg, once per day, 3 weeks). Note the abrogation of Vegfr3-Luc signal with one single administration of BO-110. Scale: p/s/cm^2^/sr (x10^6^). **(b)** Impact of the indicated administrations (in parenthesis) of BO-110 (BO, 0.8 mg/kg) on the emission of luciferase (upper panels) in xenografts of mCherry-labeled SK-Mel-103 implanted on *Vegfr3*^*Luc*^ nu/nu lymphoreporter nude mice. Same animals were also imaged for fluorescence emission to visualize mCherry (bottom panels) as a measure of tumor content. Scale, Vegfr3^Luc^: p/s/cm^2^/sr (x10^6^) and mCherry: p/s/cm^2^/sr (x10^9^). **(c)** Treatment with BO-110 of human patient derived xenografts (PDX) implanted in *Vegfr3*^*Luc*^ *nu/nu*. 42 days after implantation (when systemic luciferase was detected), animals were randomized for treatment with vehicle (V) or with 0.8 mg/kg BO-110 (BO, twice per week), and luciferase emission was acquired at the indicated times. Scale, p/s/cm^2^/sr (x10^6^). **(d)** Growth curves of *Vegfr3*^*Luc*^;*Tyr:CreERT2*;*Braf*^*V600E*^;*Pten*^*flox/flox*^ melanomas treated with BO-110 (BO, 0.8 mg/kg, 2 doses/week, 3 weeks), Vemurafenib (Vem, 50 mg/Kg, daily dose, 3 weeks), or vehicle control (V, daily dose, 3 weeks). Data correspond to average tumor area ± SD at the indicated time points. Red arrows mark the initiation of treatment (n= min 8 mice per condition). *p*=0.0007 (BO-110) and *p*=0.0353 (vemurafenib). **(e)** Growth of SK-Mel-103 implanted in *Vegfr3*^*Luc*^ *nu/nu* treated with vehicle control, 80 mg/Kg dacarbazine (DTIC, 5 doses/week, 2 weeks) or 0.8 mg/Kg BO-110 (BO, 2 doses/week, 2 weeks). Tumor volumes were measured at the indicated times and represented as average ± SD (n= min 6 mice per condition). *p*=0.0001 (BO-110) and *p*=0.0462 (DITC). **(f)** Quantification of the inhibitory effect of BO-110 (BO, 0.8 mg/kg, 2 doses/week I.P. administration, 11 weeks) on the growth of melanoma patient-derived xenografts (PDXs). Here as in panels (d) and (e), red arrows mark the initiation of treatment. *p*=0.0009. **(g)** Histological visualization of lymphatic vessel density (Lyve1 staining, brown) in representative lymph nodes of SK-Mel-103-driven xenografts in *Vegfr3*^*Luc*^ *nu/nu* mice treated with vehicle (V) or 4 doses of BO-110 (BO, 0.8 mg/kg). **(h)** Representative sentinel, axillary and brachial lymph nodes (SLN, ALN and BLN, respectively) of mCherry-SK-Mel-103-driven xenografts in *Vegfr3*^*Luc*^ *nu/nu* mice treated with vehicle (V) or 4 doses of BO-110 (BO, 0.8 mg/kg) and imaged for mCherry fluorescence to assess metastatic potential as a function of treatment. Numbers in parenthesis indicate the amount of mice with positive metastasis in at least one LN with respect to the total animals analyzed per condition. Scale for the bioluminescence images, p/s/cm^2^/sr (x10^8^).

Once references for moderate and negative anti-lymphangiogenic compounds were set, the *Vegfr*^*Luc*^ mice were interrogated for pathogen-activated molecular pattern (PAMP)-derivatives. These agents, and in particular, nanoplexes based on long synthetic dsRNA were attractive for their promising anti-metastatic activity^30–35^. Specifically, we had previously demonstrated that synthetic (poly)inosinic:polycytidylic acid, a classical mimic of viral dsRNA, can be efficiently delivered to tumor cells when packed with polyethyleneimine into bioavailable nanocomplexes^36,37^ (herein referred to as BO-110 for simplicity). BO-110 can promote an effective self-digestion of melanomas, independently of the mutational status of BRAF, NRAS and other melanoma drivers^36^. Still, it is unknown whether (and if the case how) BO-110 could impact on the development and/or maintenance of lymphovascular niches. We considered these possible functions of interest as a derivative of BO-110 (BO-112) has shown a potent antitumoral activity also in other systems, with additional roles in the immune system^38^. Moreover BO-112 and other dsRNA-based mimics are being conducted for various malignant diseases^33–35^. As shown in Figs. 1a-f, BO-110 was found as a potent inhibitor of Vegfr3-associated lymphangiogenesis, significantly exceeding the effects observed for vemurafenib and DTIC. One single administration of BO-110, although not yet sufficient to promote the collapse of tumor cells, reduced Vegfr3-luciferase emission by 80% in *Vegfr3*^*Luc*^;*Tyr:CreERT2;Braf*^*V600E*^;*Pten*^*flox/flox*^ background (Fig. 1a; see comparisons to vemurafenib). This potent reduction in Vegfr3-luciferase was also observed in *Vegfr3*^*Luc*^ mice harbouring melanoma lesions driven by aggressive human melanoma cell lines (labeled with fluorescent biomarkers to ease the visualization of tumor cells, Fig. 1b), or PDX (Fig. 1c). Importantly, 6 doses of BO-110 abrogated melanoma growth and Vegfr3 induction and melanoma growth, in both immunocompetent (Fig. S1e) and immunodeficient backgrounds (Fig. 1b, c), providing sustained therapeutic benefit in all these settings (Figs. 1d-f).

To discard unspecific effects of BO-110 on luciferase stability, we treated cells that expressed luciferase under an unrelated (SV40) promoter, and followed bioluminiscence both *in vitro* and *in vivo*. Importantly, BO-110 had no impact on luciferase emission in any of these models (Fig. S2a-d). Instead, histological staining for standard lymphatic endothelial markers (Lyve1) demonstrated an acute abrogation of tumor-driven lymphangiogenesis by BO-110 (see representative sentinel lymph nodes in Fig. 1g). Moreover, fluorescent-based imaging of tumor cells indicated that this potent anti-lymphangiogenic effect was paralleled by a striking inhibitory effect on metastatic dissemination (Fig. 1h). Importantly, in contrast to the lymphedema reported by soluble Vegfr3 traps^39^, BO-110 blocked tumor-induced neolymphangiogenesis without secondary toxicities to normal lymphatic cells (not shown). Together, these results validate *Vegfr3*^*Luc*^ reporters as a tractable platform for drug testing *in vivo*, and point to BO-110 as a distinct and selective blocker of pathogenic lymphovascular niches.

### Dual impact of BO-110 on tumor and endothelial cell-driven lymphangiogenic factors

Tumor-associated neo-lymphangiogenesis is a complex process classically induced by cytokines such as VEGFC/D (which may be secreted by aggressive cancer cells) that activate their cognate VEGFR3 receptor in lymphatic endothelial cells (LECs)^18,19^. *Vegfr3*^*Luc*^ reporters have identified a distinct mechanism whereby melanomas drive lymphatic expansion via the secretion of the heparin binding factor Midkine (MDK)^25^. Thus, we questioned which (if any) of these pathways could be modulated by BO-110. RNA-based analyses (RT-PCR) of melanoma cells treated with BO-110 showed that the expression of classical lymphangiogenic ligands *VEGFC* and *VEGFD* were either non-altered or even induced, respectively (see quantifications for two melanoma cell lines in Fig. 2a). In contrast, BO-110 repressed MDK mRNA expression (Fig. 2b and secretion (Fig. 2c) in cultured melanoma cells. Importantly, these effects were detected at early time points in which cell viability was maintained (Fig. S2e, left panels). *In vivo*, histological analyses demonstrated a potent abrogation of MDK expression in BO-110 xenografts generated by human cell lines and in PDX (see examples in Fig. 2d). Together, these data illustrate the versatility of *Vegfr3*^*Luc*^ receptors not only to identify lymphangiogenic factors such as MDK^25^, but also to uncover agents to repress them.

**Figure 2.**
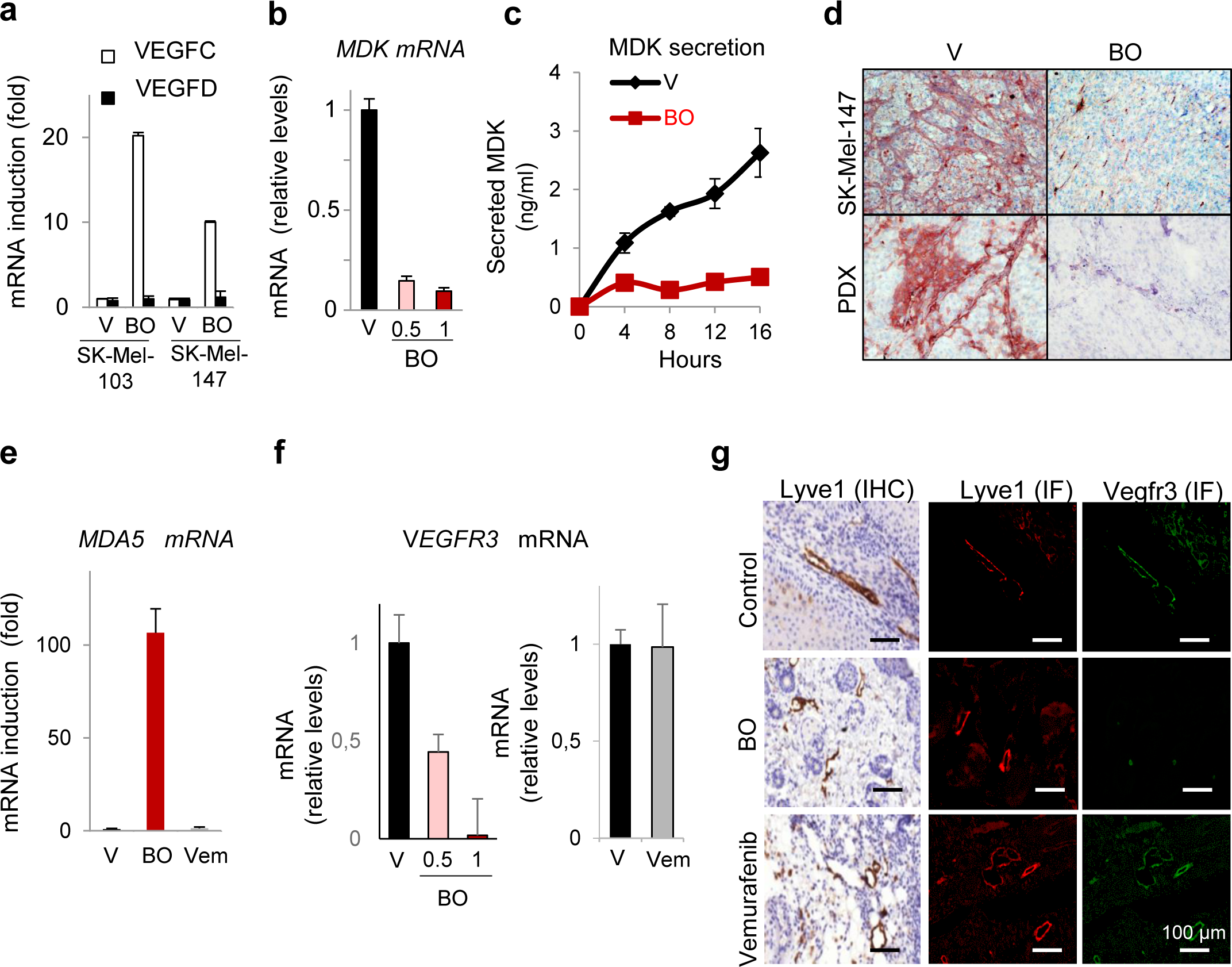
Dual transcriptional repression of *MDK* and *VEGFR3* by the dsRNA mimic BO-110. **(a)** Relative mRNA levels of *VEGFC* and *VEGFD* in the indicated melanoma cell lines 8 h after treatment with vehicle (V) or 0.5 µg/ml BO-110 (BO), as determined by qRT-PCR. Data correspond to mean ± SD of 3 experiments. **(b)** Inhibitory effect of the indicated doses of BO-110 (in µg/ml) on *MDK* mRNA expression in SK-Mel-147 determined by qRT-PCR. Data correspond to average mRNA levels of three experiments normalized to vehicle control ± SD. **(c)** Quantification of MDK secretion by ELISA in SK-Mel-147 melanoma cells treated with 0.5 µg/ml BO-110 (BO). Data correspond to mean ± SD of 3 biological replicates. **(d)** Immunohistochemical analysis of *MDK* mRNA repression (pink staining) in SK-Mel-147 xenografts and PDX lesions after treatment with BO-110 (BO, 0.8 mg/kg, 2 doses/week). Histological staining in tumors extracted from animals treated with vehicle control (V) are included as a reference. Nuclei were counterstained in with hematoxilin. **(e)** qRT-PCR analysis of relative mRNA levels of *MDA5* 16 h after treatment of HLEC with 0.5 ug/ml BO-110 (BO), 10 µM vemurafenib (Vem), or vehicle control (V). Data correspond to mean ± SD of 3 biological replicates. **(f)** qRT-PCR analysis of relative mRNA levels of *VEGFR3* 16 h after treatment of HLEC with 0.5 or 1 ug/ml BO-110 (VO), 10 µM vemurafenib (Vem), or the corresponding vehicle control (V). Data correspond to mean ± SD of 3 biological replicates. **(g)** Histological assessment of the differential impact of BO-110 (BO) and Vemurafenib (V) on neo-lymphangiogenesis in melanomas generated in *Vegfr3*^*Luc*^;*Tyr:CreERT2*; *BRAF*^*V600E*^;*Pten*^*fl/fl*^*Pten*^*flox/flox*^ mice. Panels correspond to Lyve1 detected by immunohistochemistry (IHC, brown signal) or immunofluorescence (IF, red signal), as well as to immune-detection of Vegfr3 (IF, green) performed in consecutive sections of cutaneous lesions of animals treated as in Fig. 1d.

Given the strong anti-lymphangiogenic effect of BO-110 *in vivo*, with an almost complete abrogation of luciferase signal after a single dose (Fig. 1a), we also questioned potential direct effects of this compound on the lymphatic vasculature. Analyses with cultured human lymphatic endothelial cells (HLEC) demonstrated a direct uptake of BO-110, as assessed by the induction of the cytosolic dsRNA helicase MDA5, a demonstrated sensor of this dsRNA nanoplex^36^ (Fig. 2e). Importantly, BO-110 treatment inhibited the proliferation and tubulogenic activity in three-dimensional matrices of HLECs (see below in Fig. 3d), activities known to be involved in neolymphangiogenesis^40^. To address the mechanisms whereby BO-110 affected LEC function, we next assessed whether it could directly repress *VEGFR3* levels. RT-PCR demonstrated this to be the case (Fig. 2f, left graph), without affecting the viability of cultured HLEC (Fig. S2e). This inhibitory effect of BO-110 was also found *in vivo*, by costaining for Vegfr3 and the lymphatic marker Lyve1 in BO-110 treated melanoma-bearing mice (Fig. 2g, middle panels). Interestingly, this Vegfr3-inhibitory property of BO-110 was not shared with Vemurafenib neither in cultured cells (Fig. 2f, right graph) nor in melanoma lesions *in vivo* (Fig. 2g, bottom panels). Therefore, these data further illustrate the differential impact of BO-110 and targeted therapy on lymphovascular sites. Moreover, the distinct ability of BO-110 to repress pro-lymphangiogenic factors both at the tumor and LEC level (i.e. MDK and VEGFR3, respectively) differentiates this compound from other anti-lymphangiogenic compounds in clinical testing, where the reported mode of action relies on blocking the interaction between VEGFR3 and their VEGFC/D ligands^19^, without altering VEGFR3 expression.

**Figure 3.**
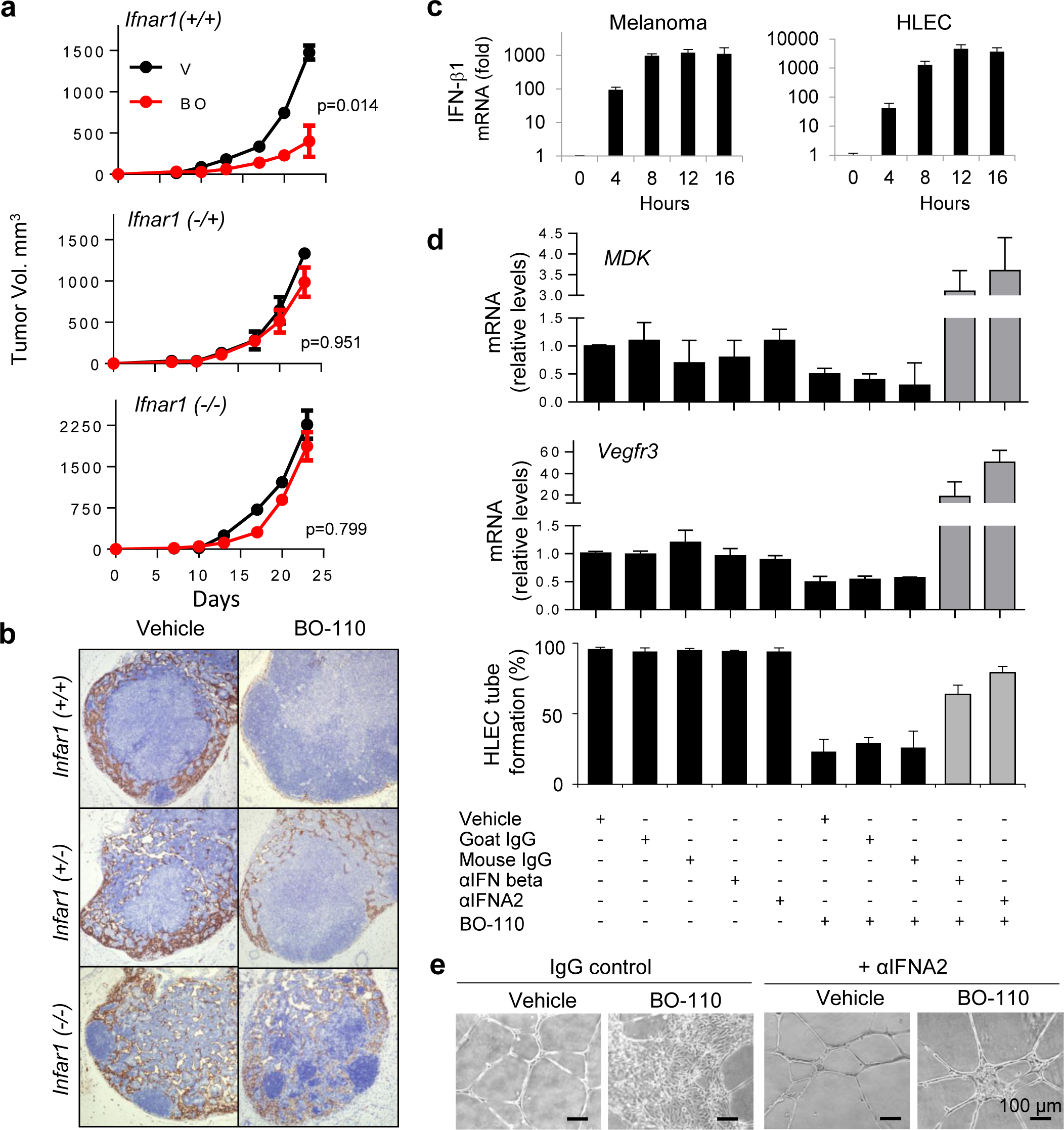
IFN-driven repressive activity of BO-110 on melanoma-induced neo-lymphangiogenesis. **(a)** B16 melanoma xenografts growth in siblings of *Infar1*^*+/+*^, *Infar1*^*+/−*^ or *Infar1*^*−/−*^ mice. Treatment started 10 days after tumor cell implantation. BO-110 was administered at 0.8 mg/kg, every third day during 2 weeks. A minimum of 6 animals were analyzed per experimental condition. Graphs show mean tumor size ± SD at each time point. **(b)** Histological analyses of lymphatic vessel density (Lyve1 staining) in representative lymph nodes of animals in (a) processed at the endpoint of the experiment (4 doses of BO-110 or vehicle control). n= min of 6 animals per experimental condition. **(c)** *IFN*β mRNA induction at the indicated times after BO-110 treatment (0.5 µg/ml) of SK-Mel-147 melanoma cells or HLEC (left and right graphs, respectively). **(d)** Quantification of the impact of BO-110 alone or in the presence of the indicated blocking antibodies for type I interferon (IFN-β or IFNA2). Upper graphs show the effect of these agents on *MDK* mRNA levels in SK-Mel-147 melanoma cell line. Similar treatments were performed on HLEC for analysis of *VEGFR3* mRNA (middle graphs) and tube formation capacity (lower graphs). Data correspond to mean ± SD of 3 biological replicates in triplicate. **(e)** BO-110-driven blockade of tube-forming capacity of HLEC and rescue with anti-IFNA2 blocking antibodies. Images correspond to cells plated in matrigel and imaged 8h after treatment with 0.5 µg/ml BO-110. See also Fig. S3c-d for additional results with anti-IFNA2 and anti-IFNβ blocking antibodies.

### Dual impact of BO-110 on tumor and endothelial cell-induced lymphangiogenic factors

Various signaling cascades have been described to upregulate MDK or VEGFR3^17–19^. However, repressors of these genes are less understood. In fact, to our knowledge, no mechanism has been reported to dually repress mRNA levels of both genes. Therefore, we focused on the identification of downstream effectors on MDA5 and other cytosolic sensors of long dsRNA, which involving transcriptional programs^41^, could act on a differential manner on tumor cells and the vasculature. Type I interferon (IFN)-related responses were attractive because they may contribute, but are not strictly required^37^ for the killing of melanoma cells by cytosolic dsRNA mimics (this function being driven by autophagy and apoptosis)^36,37^. Nevertheless, IFN plays key roles in the lymphatic vasculature as described in developmental and inflammatory processes^17^. For a first approximation to the global requirement of type I IFN on the antitumoral response of BO-110, drug response analyses were performed on mice engineered to carry mono or biallelic deletions of the IFN-α/β receptor 1 (Ifnar1)^42^. As shown in Fig. 2a, the therapeutic efficacy of BO-110 on xenografts of the isogenic melanoma cells (B16-V5) was nearly abrogated in *Ifnar1*^+/−^ and *Ifnar1*^*−/−*^ mice (see tumor growth curves in Fig. 3a). Moreover, histological staining for LYVE1 in LN of melanoma-bearing animals of the different *Ifnar1* backgrounds further revealed the requirement of type I IFN for the repressive effect of BO-110 on neo-lymphangiogenesis *in vivo* (Fig. 3b).

Next, we questioned whether IFN-α/β directly contributed to the inhibitory effect of BO-110 on melanoma cells (via MDK) and lymphatic endothelial cells (by blocking VEGFR3). Melanoma cell lines (SK-Mel-103) and HLEC where cultured and treated independently to dissociate roles of IFN in both cell types. Transcriptomic profiles extracted from cDNA arrays in BO-110 treated SK-Mel-103 cells (GSE14445)^36^ demonstrated a classical IFN-associated response (see Fig. S3a). Moreover, qRT-PCR based analyses confirmed a 100-1000 fold induction of *IFNβ1* in SK-Mel-103 as early as 4-8h after treatment with BO-110 (Fig. 3c, left panels) paralleling the inhibition of MDK (Fig. 2c). This IFN upregulation was functionally relevant, as blocking antibodies against IFNβ or against IFNα/β Receptor Chain 2 (IFNA2) counteracted the impact of BO-110 on *MDK* expression (Fig. 3d). Regarding HLEC, and consistent with these cells uptaking BO-110 (as reflected by MDA5 activation; Fig. 2e), type I IFN (particularly IFNβ) was significantly upregulated by this compound (Fig. S3b), also at very early time points after treatment (Fig. 3c, right panels). Importantly, IFNβ or IFNA2-blocking antibodies also rescued the repression of BO-110 on VEGFR3, allowing for an efficient proliferative (Fig. S3c) and tubulogenic activity of HLEC cells (Fig. 3d,e; see additional controls in Fig. S3d). Together, these results identify a dual and selective mode of action of BO-110 repressing *MDK* in melanoma cells and *VEGFR3* in LEC. The identification of Type I IFN as a common driver of both processes opens new avenues of research on how to deactivate tumor-driven lymphovascular niches.

### Long-term anti-metastatic effects of BO-110 *in vivo* prevent metastatic relapse after surgery

Collectively, the data above illustrate how the *Vegfr3*^*Luc*^ mice can be geared to the discovery of anti-lymphangiogenic factors with novel modes of action. We envision this information to be relevant for melanoma patients that appear with metastatic disease at diagnosis. Still, an additional important need in the field is to monitor (and attack) metastatic relapse after surgical excision of primary lesions. Therefore, the *Vegfr3*^*Luc*^ mice were interrogated for their capacity to mimic this scenario, and thus, to serve to screen for adjuvant treatments. To this end, xenografts of human mCherry-labeled SK-Mel-147 melanoma cells were implanted subcutaneously, for surgical excision of the cutaneous lesions once systemic VEGFR3-associated bioluminescence was identified. As shown in Fig. 4a, surgery resulted in a progressive reduction in systemic Vegfr3-luc emission, indicating the requirement of tumor-derived signals to maintain distal neo-lymphangiogenesis. However, and as in the case for highly aggressive melanomas in the clinic, animals ultimately succumbed to relapsed metastases, revealed by live imaging of Vegfr3-bioluminiscence (Fig. 4b, right panels), and monitoring of tumor cell burden by mCherry-fluorescence (Fig. S4). Importantly, BO-110 (4 doses every third day, starting 4 days after surgery) prevented both the re-acquisition of lymphangiogenesis and the subsequent tumor relapse (Fig. 4a bottom panels). Importantly, the efficacy of BO-110 was long-lasting, as 90% of animals remained tumor-free 8 months after treatment (see Kaplan Meier survival curves in Fig. 4b; p=0.0001). Together, these results emphasize the versatility of *Vegfr3*^*Luc*^ GEMM reporter mice for non-invasive studies of melanoma initiation and progression, as well as a tractable platform for pharmacological screening for actionable targets to prevent and attack tumor metastasis.

**Fig. 4.**
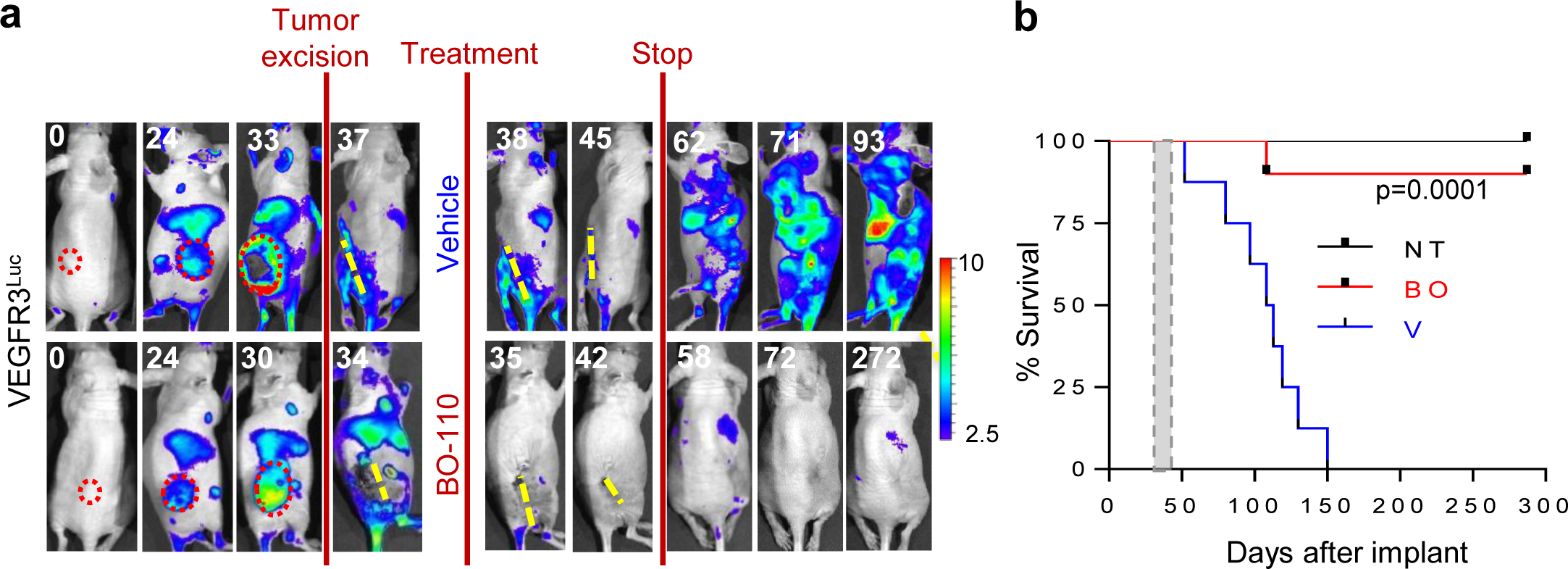
Long-term impact of BO-110 preventing metastatic relapse and extending half life of treated mice. **(a)** Efficacy of BO-110 as adjuvant (preventing relapse after surgical removal of the primary lesion). Shown are representative images of *Vegfr3*^*Luc*^ mice implanted with mCherry-SK-Mel-147 and imaged for luciferase emission prior and after tumor removal. Animals were left to recover from surgery (4 days), and then treated for 2 weeks (4 doses) with 0.8mg/Kg BO-110 or vehicle control (n= 8 for control and n=10 for treatment arm). Scale, p/s/cm^2^/sr x10^6^. See Fig. S4 for the corresponding mCherry signal (to visualize tumor burden) prior and post-surgery, and at the indicated time points after treatment. **(b)** Kaplan-Meier survival curves of animals treated as in (f). 8/8 animals treated with vehicle (V) control had to be sacrificed for humane reasons 110 days after surgery. 9/10 animals in the BO-110 arm (V) remained tumor-free 8 months after stopping treatment. The grey box marks the period of treatment with BO-110.

## DISCUSSION

Drug testing and target validation in melanoma are compromised by the lack of physiological systems for non-invasive imaging of metastasis. Here we exploited immunocompetent and immunodeficient *Vegfr3*^*Luc*^ knock-in mice as an *in vivo* screening platform for anticancer agents. Specifically, our data support these mice as a versatile resource for spatio-temporal analyses of drug response on established tumors, as well as to test for compounds able to prevent tumor formation prior or after surgical excision of cutaneous lesions. Specifically, we demonstrate the efficacy of these models with the identification of potent anti-lymphangiogenic functions of the dsRNA mimic BO-110 (see model in Fig. 5). Guided by the visualization of an acute blockade of lymphangiogenesis at primary and distal sites, a dual IFN-dependent repressive function of BO-110 was found on tumor cells (inhibiting MDK expression and secretion) and on lymphatic endothelial cells (acting on *Vegfr3* mRNA levels). These functions, distinct from the reported action of lymphangiogenic factors in clinical testing^19^, were not recapitulated by FDA-approved therapies (BRAF inhibition and treatment with alkylating agents). Of note, BO-110 did not promote detectable damaging effects on normal lymphatic vessels. Therefore, the *Vegfr3*^*Luc*^ reporters have uncovered differential effects of dsRNA mimics not only in cancer cells and their associated vasculature, but on pathological versus normal lymphatic vessels as well.

**Fig. 5.**
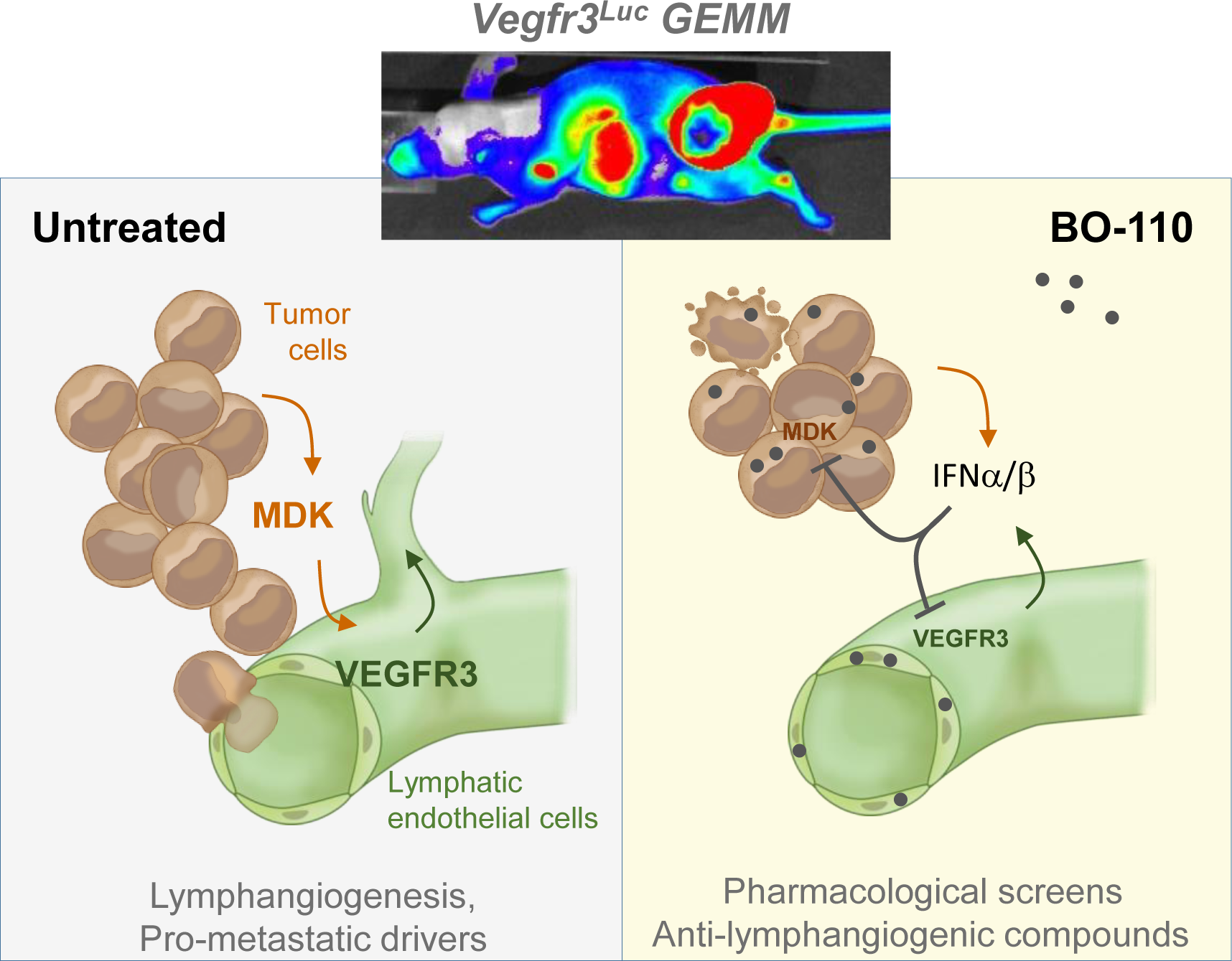
Model summarizing the main conclusions of this study. Versatility of the *Vegfr3*^*Luc*^ “MetAlert” lymphoreporter mice for gene discovery and pharmacological testing of anticancer agents. We had previously reported that non-invasive imaging of *Vegfr3-*associated neo-lymphangiogenesis allowed for the identification of MDK as a driver of Vegfr3-induced sprouting and expansion of the lymphangiogenic vasculature, a process activated at early stages of melanoma development^25^. Here we show that the MetAlert mice can represent a cost-effective strategy to screen for inhibitors of lymphangiogenesis and define their therapeutic efficacy in immunocompetent and immunodeficient settings (GEMM models). Using these MetAlert mice, we identified a novel role of the dsRNA mimic BO-110 as an effective blocker of neo-lymphangiogenesis. Mechanistically, BO-110 was found to represent an unexpected dual repressor of *MDK* and VEGFR3 transcription, an effect mediated by type I IFN-signaling. Of note, the ability of the MetAlert mice to monitor premetastatic niches after surgery allow also for surrogate analyses of neo-adjuvant therapies (here also shown to be effective for BO-110 treatments).

Together, these data have implications at various levels. In particular, this work illustrates the impact of treatments that target both cancer cells and distal lymphovascular pre-metastatic niches (as the latter may be activated very early in the metastatic process). In this context, despite recent great therapeutic advances, the melanoma field is still in need of compounds, particularly in adjuvant settings, that provide long-term responses that are not restricted to specific patient subsets^28,43^. Pre-clinical studies support BO-110 and other formulations of long synthetic dsRNA as potent therapeutic agents for their ability to kill melanoma cells otherwise resistant to standard therapies^32,36–38^. We had previously linked these effects to an IFN-independent activation of apoptosis and autophagy in tumor cells^36,44^. The *Vegfr3*^*Luc*^ mice revealed now an additional level of action of dsRNA mimics as potent anti-lymphangiogenic agents. This is relevant as clinical trials with dsRNA mimics are underway in various pathologies^33–35,38^. Moreover, finding a strategy to block VEGFR3 transcription may also be attractive from a pharmacological perspective, as to date, anti-lymphangiogenic factors in clinical trials (tyrosine kinase inhibitors, VEGFC/D traps or VEGFR3-VEGFC/D interaction competitors) act by interfering with VEGFR3 function, not the expression of this protein^19^.

Here we addressed BO-110 and FDA-approved drugs as a single agents, but the *Vegfr3*^*Luc*^ mice could represent a cost-effective strategy for the analyses of drug combinations. For example, the induction of IFN by BO-110 is relevant as this signaling cascade can exert multiple systemic effects on the innate immune system^31,45^. These can include for example the mobilization of dendritic, myeloid, NK and T cells, that may favor the actively pursued effects of immune checkpoint inhibitors^32^. Finally, it should be noted that although this study focused on melanoma, the *Vegfr3*^*Luc*^ mice could be exploited for gene discovery and pharmacological analyses in other cancer types. The ability to compare immunocompetent and immunosuppressed backgrounds, add yet further physiological relevance to whole-body imaging of lymphangiogenesis in tumor progression and therapeutic response.

## METHODS

### Mouse breeding, induction of nevi and melanomas in Vegfr3^Luc^ GEMM, and drug treatments in vivo

The *Vegfr3*^*Luc*^ nu/nu immunodeficient mice and the *Vegfr3*^*Luc*^; *Tyr:CreERT2;Braf*^*V600E*^; *Pten*^*flox/flox*^ animals were generated as described in Olmeda et al. (accompanying paper). Specifically, strains used in this study are as follows: *Vegfr3*^*neoEGFPLuc*^ (Flt4^tm1.1Sgo^), initially engineered in *CD-1, C57BL6/J-albino* background^26^; *nu/nu* (*Crl:NU(Ico)Foxn1nu*); *Tyr::CreERT2/1Lru*^46^; *Braf*^*CA*^ (*Braf*^*tm1Mmcm*^)^27^ and *Pten*^*tm2Mak*^ ^47^. Melanomas in the *Tyr::CreERT2* strains were induced in 14 week-old mice by topical treatment with 5 µl of 5 mM 4-hydroxytamoxifen. BO-110 (pIC-PEI) was prepared as described before^36^. When indicated, 0.8 mg/Kg BO-110 was injected intravenously every three days (for a total of 4 administrations, unless indicated otherwise). DTIC (80 mg/kg; Sigma Aldrich) was administered intraperitoneally in cycles of 5 consecutive days per week^48^. Vemurafenib (Selleck Chemicals, TX) was prepared and administered at 50 mg/Kg (orally, once a day, during 3 weeks) as previously described^49^.

### Non-invasive imaging of tumor growth and neo-lymphangiogenesis in vivo

Non-invasive imaging of luciferase in the *Vegfr3*^*Luc*^ GEMM was performed using an IVIS-SPECTRUM imaging system (Perkin Helmer, Baesweiler Germany). Animals were anesthetized with isoflurane and injected intraperitoneally with 150 mg/kg luciferin (Perkin Helmer). Sequential images were obtained after luciferin injection and the maximum light emission was determined for each animal as previously described^26^. Photons emitted from specific regions were quantified using Living Image software (Caliper Life Sciences). *In vivo* luciferase activity is presented in photons per second per square centimeter per steradian (radiance). All experiments with mice were performed in accordance with protocols approved by the Institutional Ethics Committee of the CNIO and the Instituto de Salud Carlos III.

### Melanoma cell xenografts, patient-derived xenografts and spontaneous metastasis assays

Xenografts of established melanoma cell lines were generated in 14-week old female *Vegfr3^Luc^ nu/nu* mice by subcutaneous implantation of 1 × 10^6^ cells. Tumor growth was recorded every two days by measuring the two orthogonal external diameters using a calliper. Tumour volume was calculated using the formula (a x b^2^ x 0.52). Tumors were excised and processed for histological analysis when they reached 1.5 cm^3^. For spontaneous metastasis assays, SK-Mel-147 tumors were grown in the same conditions as described above until they reached a size of 1.2 cm^3^. Surgical excision was then performed under analgesics (buprenorphine 0.05 mg/day), and animals were left to recover for subsequent image of tumor growth and luciferase imaging at increasing time periods. When indicated, treatments for prevention of metastatic relapse were initiated 4 days after surgery.

Patient derived xenografts (PDX) were generated from biopsies of skin metastasis obtained from the Hospital 12 Octubre, Madrid, under their appropriate ethical protocols and provided to the investigators as anonymized lesions. These biopsies were excised into 4 mm cubes, embedded in Matrigel (BD) and implanted in the back of highly immunodeficient NSG mice (NOD.Cg-*Prkdc*^*scid*^ *Il2rg*^*tm1Wjl*^/SzJ). Once the tumors reached 1000 mm^3^ they were excised, processed again in 4 mm cubes and re-implanted in the back of 3-6 NSG mice for amplification and subsequent reimplantation in *Vegfr3*^*Luc*^ *nu/nu*.

### Histological analyses of gene expression in mouse tumors

Histological analyses of tissue architecture and expression of lymphangiogenic markers were performed on biopsies fixed in formalin and embedded in paraffin. Sections were prepared for hematoxylin-and-eosin (H&E) staining. For immunostaining, 3 µm paraffin sections were deparaffinized and placed in PBS. Slides were incubated with the indicated primary antibodies as described below and developed with Ultravision ONE Detection System kit (Thermo Scientific. TL-015-HAJ) using Permanent Mounting Medium (Prolong, Thermo Scientific).

For immunofluorescence, tissue sections were deparaffinized, incubated overnight with primary antibodies at 4 °C in a humidified chamber and then rinsed and incubated with fluorescent secondary antibodies for 1 hour at room temperature. Nuclei were counterstained with Prolong Gold + DAPI (Invitrogen, concentration 5 µg/mL).

Antibodies were used as follows:

| Antibody | Code | Species | Dilution | Provider |
| --- | --- | --- | --- | --- |
| mVegfr3 | AF743 | Goat | 1:25 | R&D Systems |
| mLyve-1 | ab14917 | Rabbit | 1:500 | Abcam |
| Trp2 | sc-10451 | Goat | 1:500 | Santa Cruz |
| p-ERK | 9101 | Rabbit | 1:200 | CELL Signaling Technology |

### Quantitative reverse transcription-PCR

RNA purification from melanoma tissue samples and real-time reverse transcription-PCR (qRT-PCR) were performed essentially as described^36^ using the following primers:

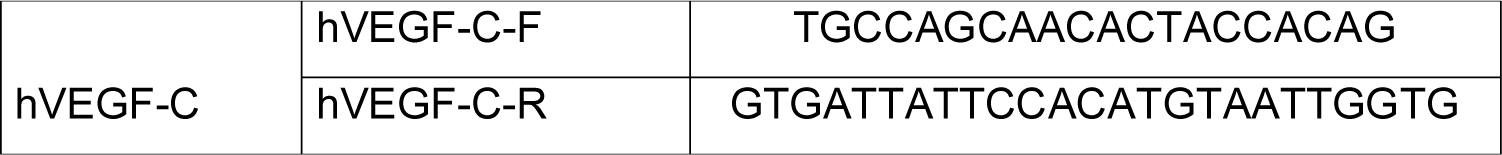
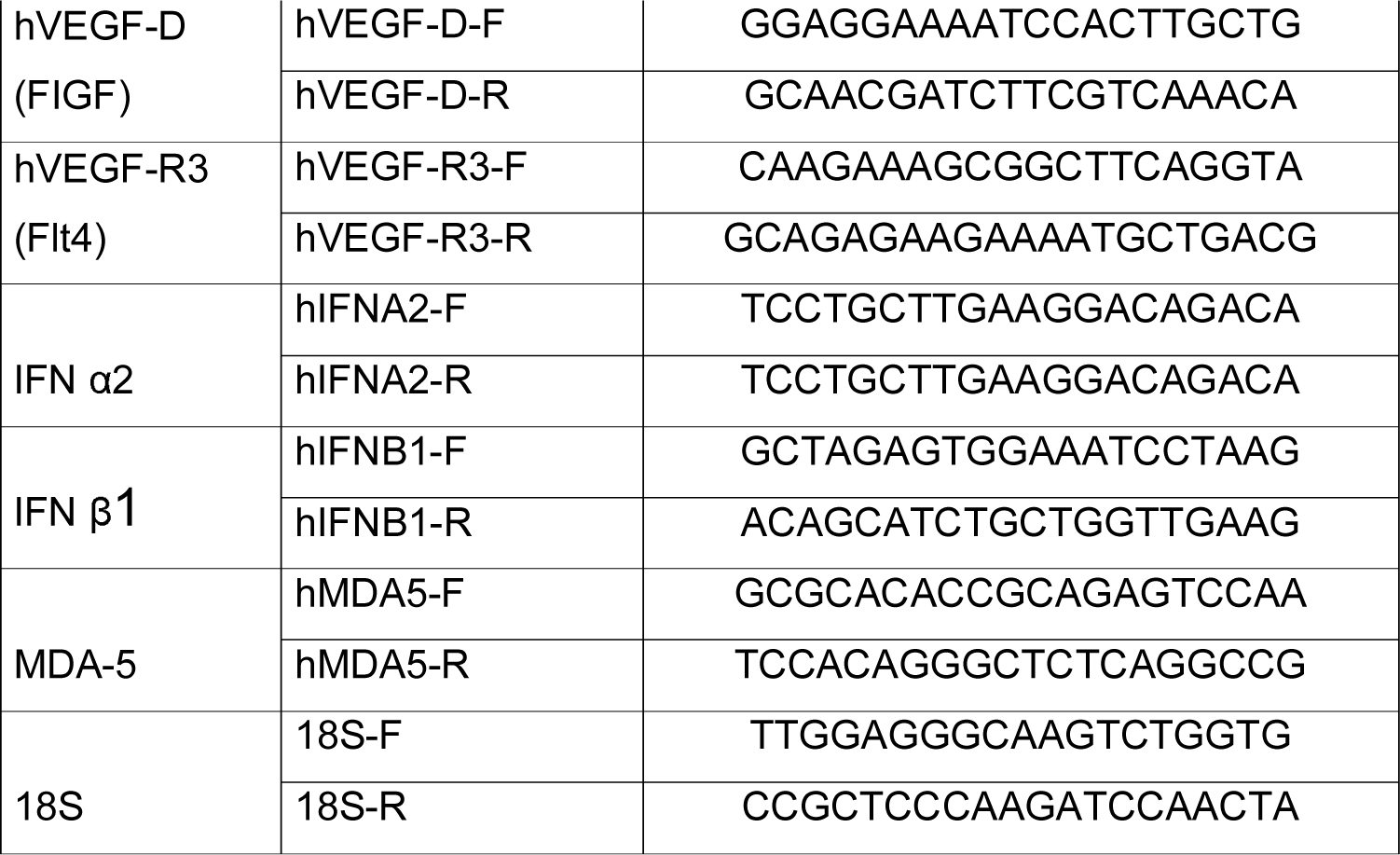

### Functional analyses in Human Lymphatic Endothelial Cells (HLEC)

Normal Human Lymphatic Microvascular Endothelial Cells (HMVEC-dLy) (Lonza, MD) referred in the text as HLEC, were grown as tissue culture monolayers (2-D) in EGM™-2 medium supplemented with MV BulletKit (Lonza). For tube-formation assays, three-dimensional cultures (3-D) were prepared with 2.5 × 10^5^ cells seeded in MW6 plates covered by a layer of matrigel (BD, NJ). When indicated, the cells were pre-treated for 12h with 0.5 µg/ml BO-110 or vehicle control. Treatment was maintained thereafter. Pictures were acquired 12 hr after seeding the cells.

### Type I IFN blocking assays

For type I interferon blocking assays, human IFN-β blocking antibody (Clone AF814; R&D) and its corresponding isotype (Goat IgG, R&D Systems) were used at a concentration of 0.2 µg/ml. Anti-Interferon-α/β Receptor Chain 2 Antibody (IFNA2, clone MAB1155; Millipore) and its corresponding isotype (Mouse IgG2a, clone GC270; Millipore) were used at 0.1 µg/ml as suggested by the provider. Where indicated, these reagents were added simultaneously with BO-110 to the culture media.

## Abbreviations

GEMM: Genetically engineered mouse models
HLEC: human lymphatic endothelial cells
LN: lymph node
PDX: patient-derived xenografts
BO-110: bioavailable nanoplexes of (poly)inosinic:polycytidylic acid packed with polyethyleneimine

## ACKNOWLEDGEMENTS

The authors thank all the colleagues in the CNIO Melanoma Group, particularly, Lisa Osterloh for help and support; José A Esteban (CSIC-UAM) for critical reading of this manuscript; Lionel Larue (INSERM; France) and Martin McMahon (Hunstman Cancer Center, USA) for the Tyr:CreERT2 and Braf^CA^ mouse strains, respectively; and Ignacio Melero at Hospital Clinico, Pamplona, Spain for Ifnar1 deficient mice. Isabel Blanco, Soraya Ruiz, Virginia Granda, Antonio Camara and Alba de Martino (CNIO) for technical assistance. M.S.S. is funded by grants from the Spanish Ministry of Economy and Innovation (SAF2014-56868-R; SAF2017-89533-R), the Asociación Española Contra el Cáncer (AECC), TV’13-20131430 (Marato de TV3), the Worldwide Cancer Research, an Established Investigator Award by the Melanoma Research Alliance (MRA), and a L’Oreal-Paris USA-MRA Team Science Award for Woman in Scientific Research. The CNIO Proteomics Unit belongs to ProteoRed, PRB2-ISCIII, supported by grant PT13/0001. J.M. is also supported by Ramon y Cajal Programme (MINECO) RYC-2012-10651.

## AUTHOR CONTRIBUTIONS

M.S.S. and D.O. conceived and designed all the studies in this work. S.O conceived and developed the *Vegfr3*^*EGFPLuc*^ (*Flt4*^*tm1.1Sgo*^) and *Vegfr3*^*Luc*^ *nu/nu* mouse models, contributed to experimental design, discussed data and revised the manuscript. D.T and M.S.S. developed BO-110, and D.T. contributed to the experimental design of the BO-110 treatments. D.O. developed the protocols for the analysis of tumor-induced lymphangiogenesis *in vivo* (pre and post-surgery, and pre-post-treatment with BO-110), and was in charge of the histological assessment of Vegfr3, Lyve1, and CD31. D.C contributed to the anti-melanoma drug treatment experiments and performed the treatments on *Ifnar1* deficient models; she helped with data analysis of the *in vivo* experiments and revised the manuscript. E.R-F and P.O.R. and J.L.R-P provided fresh melanoma biopsies for the generation of PDX. N.I. and J.M. contributed to data analysis. T.G.C. and E.C. were in charge of animal breeding and genotyping and provided technical assistance. D.A-C contributed with technical assistance. The manuscript was written by M.S.S and D.O., revised by S.O and D.A-C. and approved by all authors. M.S.S. supervised the project.

## AUTHOR INFORMATION

### Conflict of Interest

María S. Soengas and Damia Tormo are co-founders of BiOncotech Therapeutics, a small biotechnology company interested in the development of dsRNA-based treatments for aggressive cancers.

---

**Figure S1.**
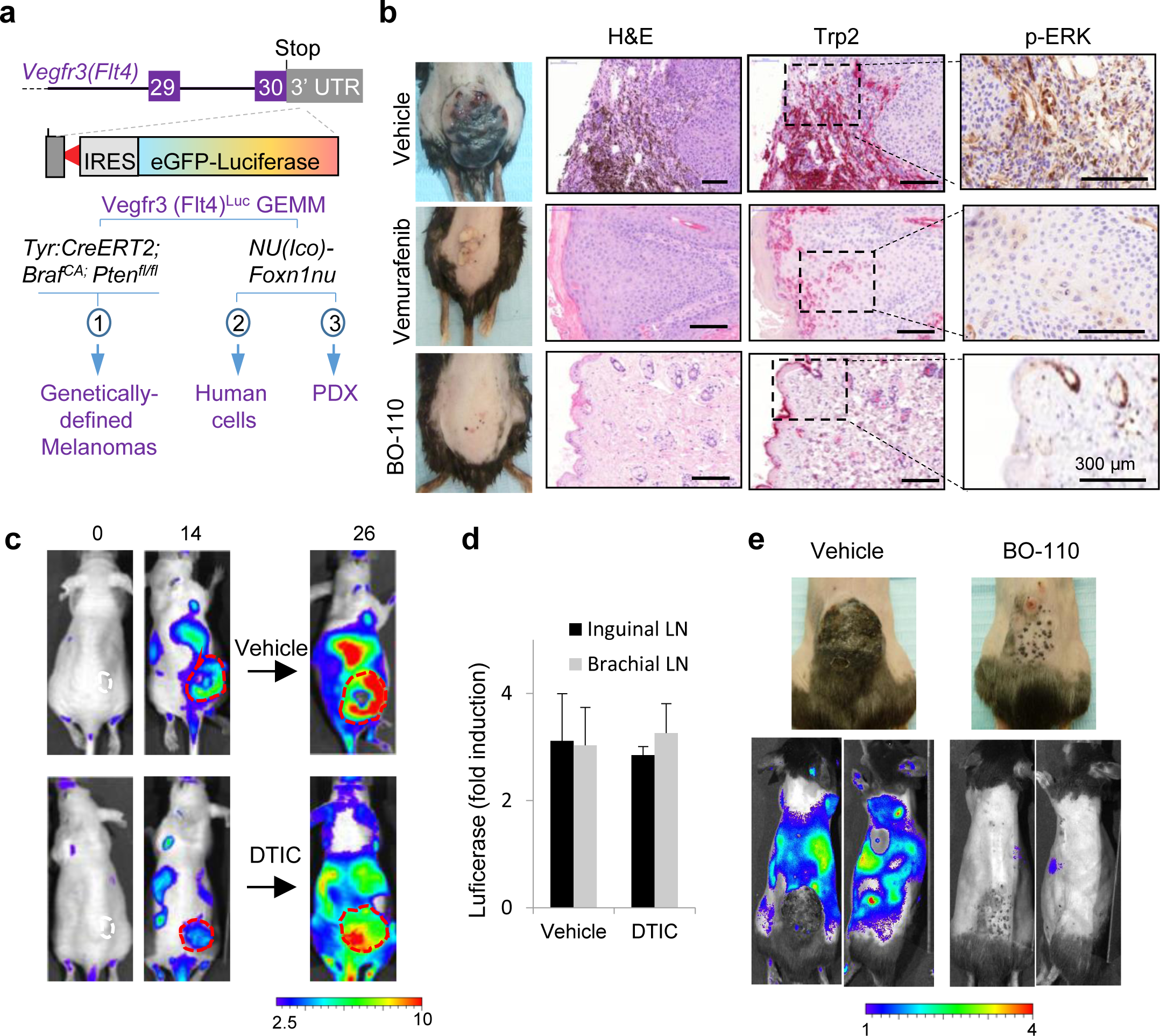
Anti-lymphangiogenic activity of BO-110 not shared by FDA-approved antimelanoma agents. **(a)** Schematic representation of the *Vegfr3*^*Luc*^ genetically engineered mouse models (GEMM) used in this study to assess melanomas driven by melanocytic-specific induction of oncogenic *Braf*^*V600E*^ in a *Pten* deficient background (1), as well as to monitor xenografts of human cells (2) and patient-derived specimens (PDX, 3). **(b)** Cutaneous melanomas generated in *Tyr:CreERT2; Braf*^*V600E*^; *Pten*^*flox/flox*^ mice treated with vehicle, BO-110 (0.8 mg/kg, twice a week during 3 weeks), or Vemurafenib (50 mg/kg, daily, 3 weeks). Left panels correspond to representative animals at the endpoints of treatment. Histological sections of these lesions are shown in the right panels, stained for H&E to visualize tissue architecture, or for Trp2 (pink) to label melanocytic cells. Insets (dotted squares) were magnified for visualization of phosphorylated ERK (p-ERK). Scale bars, 300 µm. **(c)** Tumor growth and Vegfr3-driven luciferase emission by SK-Mel-103 implanted in *Vegfr3*^*Luc*^ *nu/nu*. Tumors were left to growth until luciferase emission was clearly detectable for subsequent systemic treatment of animals with vehicle control or 80 mg/Kg DTIC. Numbers on the left correspond to days after tumor cell inoculation. Red doted lines marks tumor area. Scale, p/s/cm^2^/sr (x10^6^). **(d)** Quantification of the Vegfr3-driven emission in sentinel (inguinal) and distant (brachial) lymph nodes of animals in **(c). (e)** Impact of BO-110 on *Tyr:CreERT2*; *BRAF*^*V600E*^;*Pten*^*flox/flox*^; *Vegfr3*^*Luc*^ mice. Upper panels correspond optical photographs of animals treated with vehicle or with 6 doses of BO-110 (0.8 mg/kg, twice a week, 3 weeks), and depilated to ease in the imaging. These same animals are shown in the bottom panels for luciferase emission centering on the tumor or on sentinel LN. Scale, p/s/cm^2^/sr (x10^6^).

**Figure S2.**
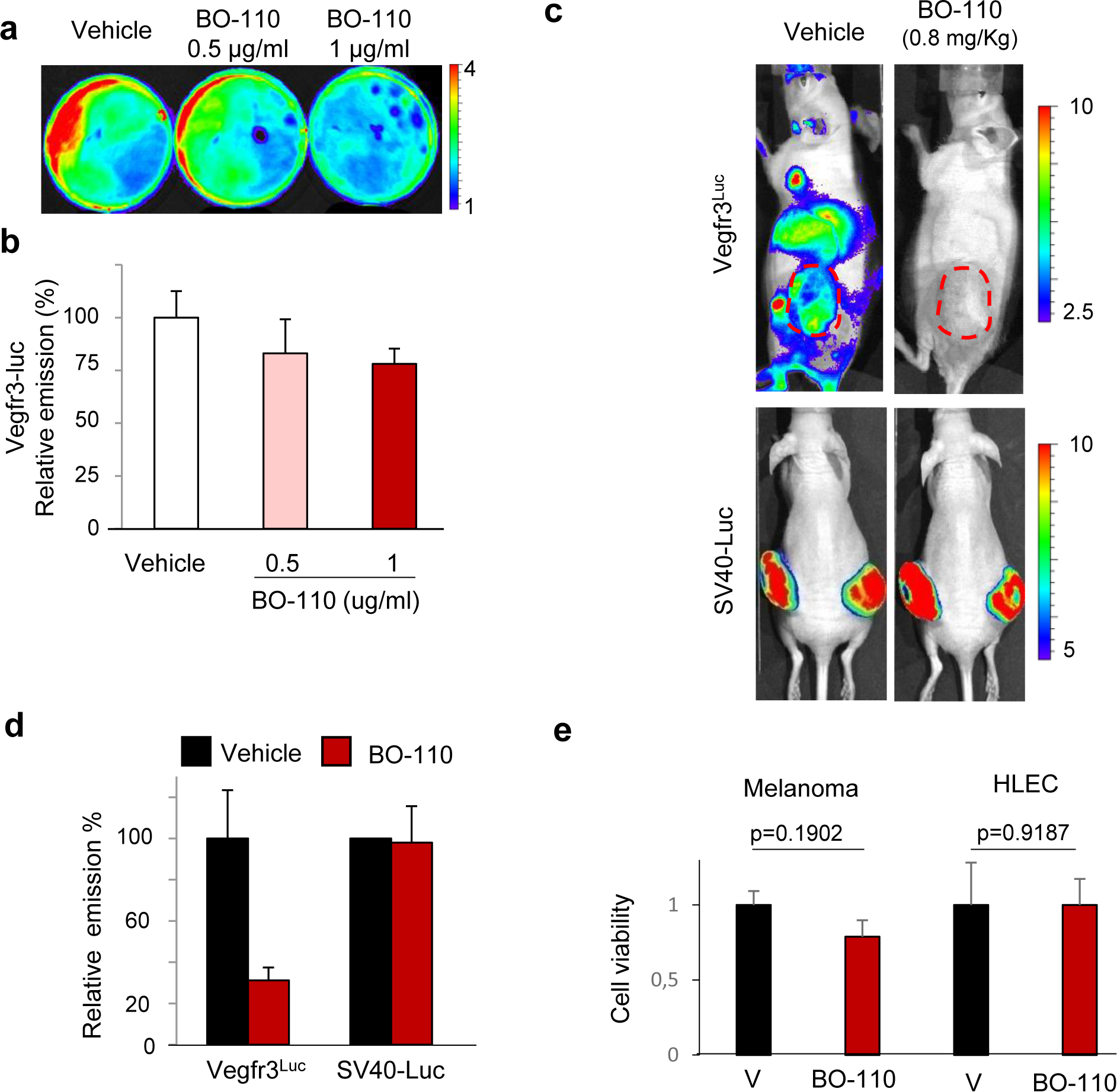
Controls for BO-110 driven inhibition of *Vegfr3*. **(a)** Lack of significant inhibitory effect of BO-110 on Luciferase emission driven by the *SV40* promoter. Shown are 3 cm plates containing an equivalent number (2 × 10^6^ cells) of SK-Mel-103 transduced with *SV40-Luc* reporter construct, and treated as indicated for 12 h. Scale: x10^7^ p/s/cm^2^/sr. Average emission signal was represented in (**b)** with respect to the untreated controls. Error bars correspond to SD of 3 experiments. **(c)** Comparative impact of BO-110 (0.8 mg/kg) on luciferase emission driven from the Vegfr3^Luc^ knock in, or from an unrelated promoter (SV40-Luc). To this end, SK-Mel-103 was divided into two populations: (i) to be transduced with an empty vector and implanted in Vegfr3^Luc^ nu/nu; and (ii) to be transduced with SV40-Luc for subsequent implantation in control mice. Scale, p/s/cm^2^/sr (x 10^6^). **(d)** Quantification of luciferase emission in the indicated animal groups treated as in (c). While Vegfr3^Luc^ signal was abrogated by BO-110, this was not the case for the luciferase driven by SV40 promoter. **(e)** Maintenance of the viability of SK-Mel-147 and HLEC estimated by trypan blue exclusion assays 16h after treatment with BO-110.

**Figure S3.**
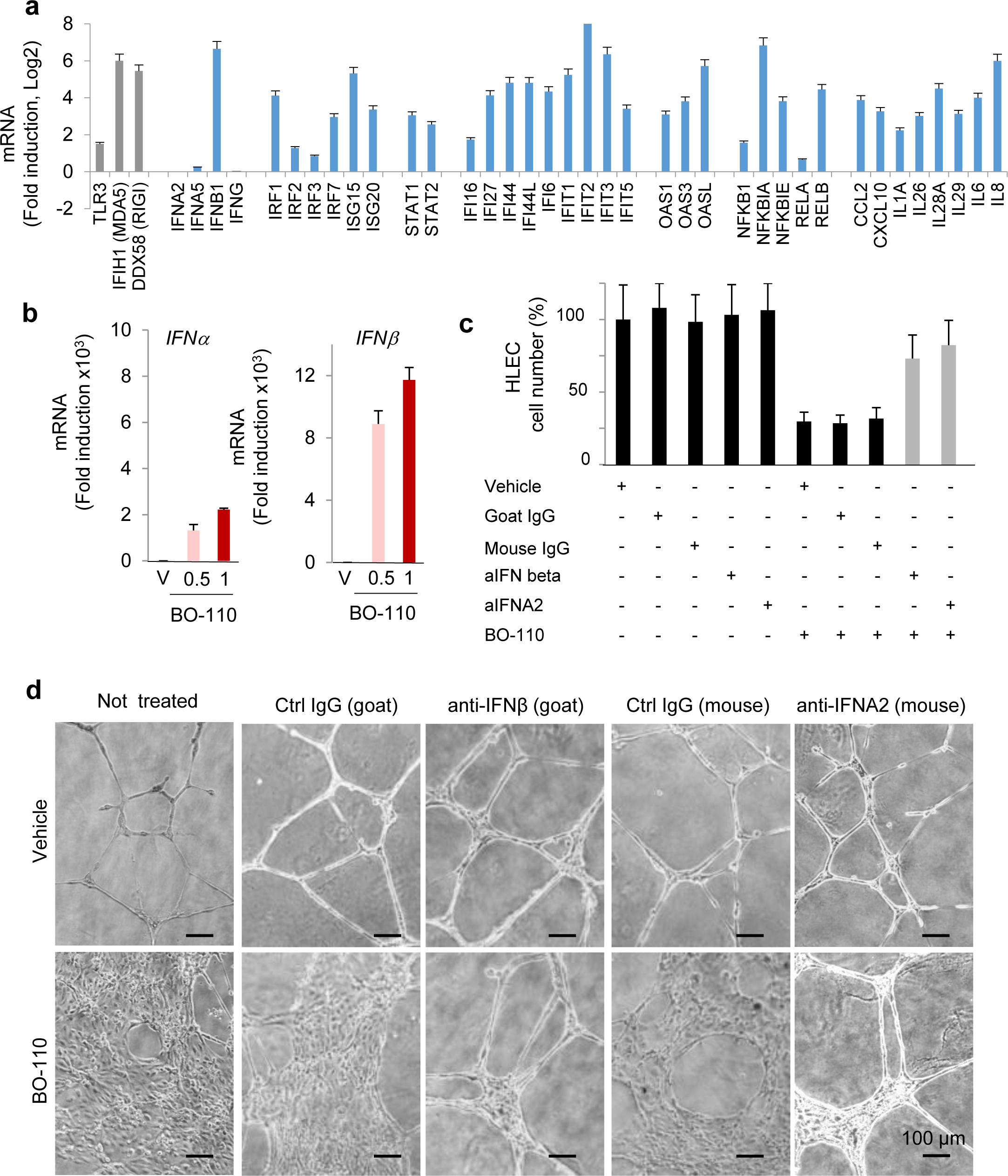
BO-110 blocks lymphangiogenesis in a type I IFN-depending manner. **(a)** Expression profile analysis of changes in the mRNA levels of the indicated IFN-responsive genes determined 10h after BO-110 treatment of SK-Mel-147. MDA5, the dsRNA sensor of BO-110, is shown as a reference (grey bar). Data correspond to triplicates analyzed by cDNA arrays. **(b)** Type I *IFN* mRNA induction (*IFNα* and *IFNβ*) in HLEC cells treated for16h with the indicated amounts of BO-110. **(c)** BO-110-driven blockade of HLEC proliferation and rescue with blocking antibodies for anti-IFNβ (goat) and anti-IFNA2 (mouse). Controls with anti-IgG from goat or mouse sources are included as a reference. Data correspond to mean ± SD of 3 biological replicates. **(d)** Representative images of the inhibition by BO-110 of the tubulogenic capacity of HLEC rescued by Type I IFN blocking antibodies (anti-IFNβ and anti-INFA2). Bars 100 µm.

**Figure S4.**
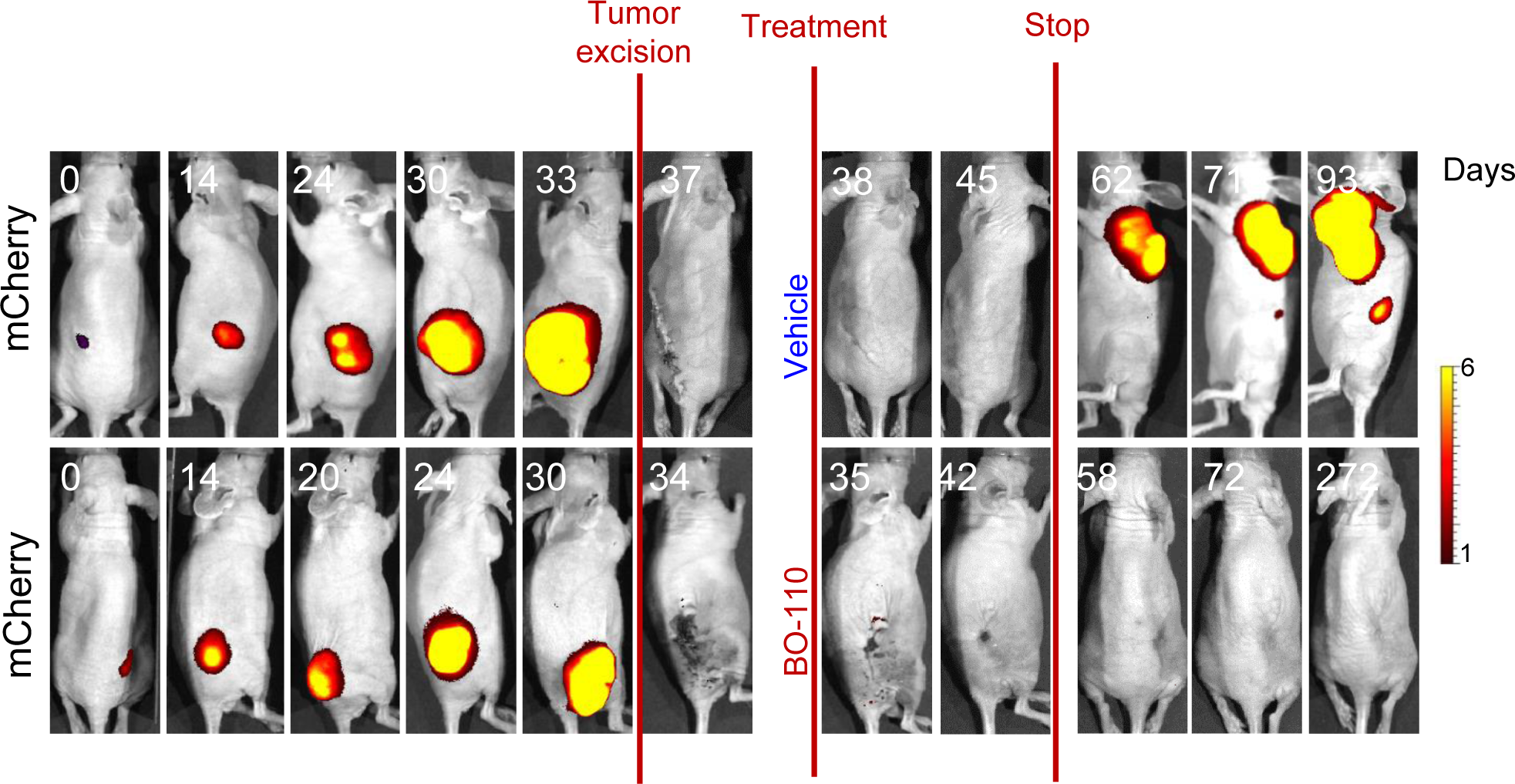
Efficacy of B0-110 as an adjuvant preventing lymphangiogenesis and metastatic relapse after surgery. Shown are representative images of *Vegfr3*^*Luc*^ mice implanted with mCherry-labeled SK-Mel-147 for fluorescence-based monitoring of tumor growth and relapse after the excision of the cutaneous lesions. Animals were left to recover from surgery (4 days), and then were treated for 2 weeks (4 doses) with 0.8mg/kg BO-110 (n=10 animals) or vehicle control (n=8 animals). Animals continued to be imaged at different time points after stopping treatments. Note the tumor relapse (fluorescence signal) in vehicle-treated animals that is prevented by BO-110. Vegfr3Luc emission and survival curves are shown in Fig. 4a, b, respectively. Scale, p/s/cm^2^/sr (x10^6^).

